# Nidogen/NID-1 guides regenerating motor axons in the mature nervous system

**DOI:** 10.64898/2026.03.13.711544

**Authors:** Eli Min, Wenjia Huang, Alexandra B Byrne

## Abstract

Restoring function to injured axons requires not only regeneration, but also accurate guidance and synapse reformation. Our understanding of how regenerating axons navigate the mature nervous system remains limited, as growth cones confront a cellular and molecular landscape distinct from development. Using *Caenorhabditis elegans*, we found that the basement membrane protein Nidogen (NID-1) promotes local guidance of regenerating motor axons in the mature nervous system by facilitating their growth alongside neighboring intact neuronal processes. Regenerating cholinergic axons preferentially track the branched dendrites of the PVD mechanosensory neuron or, in the absence of PVD dendrites, are guided alongside GABAergic commissures instead. Loss of *nid-1* disrupts this guidance, reducing axon–PVD colocalization, increasing displacement from the pre-injury point of contact with the dorsal nerve cord, and disrupting synapse reformation and functional recovery. Tissue-specific rescue indicates that NID-1 expressed by body wall muscles or the hypodermis is sufficient to guide regenerating axons, whereas muscle-derived NID-1 is required to restore synapse reformation. Genetic data indicate that NID-1 function guides regenerating axons in coordination with laminin and integrin. Although this complex primarily directs cholinergic motor axons, ectopic integrin expression in GABAergic neurons is sufficient to reroute their regenerating axons alongside PVD dendrites in a NID-1-dependent manner. Together, these findings identify a NID-1-dependent post-developmental mechanism for directing regenerating axons and promoting functional repair.

## Introduction

Functional axon regeneration requires that an injured neuron regrow to the correct location and re-establish synapses with the appropriate cells. In the central nervous system, axon regeneration is actively inhibited after injury, thereby blocking functional restoration^1–4^. Manipulating key inhibitors of axon regeneration, for example, deletion of phosphatase/tensin homolog (PTEN) and suppression of cytokine signaling 3 (SOCS3) induces robust regeneration of injured retinal ganglion cells in the optic nerve^5–8^. However, even when axons are capable of regenerating, they often display severe guidance defects and fail to reach their appropriate targets^9–11^. Consequently, improving axon regeneration alone is not sufficient to achieve pre-injury levels of recovery after nerve injury. In the developing nervous system, axon guidance is intricately regulated, both spatially and temporally. These developmental guidance cues cannot simply be co-opted to guide regenerating axons in the adult nervous system, where the molecular landscape and extracellular matrix are significantly different. While many axon guidance cues are continuously expressed into adulthood, only some maintain their ability to guide growth cones, others acquire the opposite function in the mature nervous system^12^, and regenerating growth cones must navigate a remodeled and more complex extracellular space^12–15^. The molecular and cellular mechanisms that guide regenerating axons through this adult environment remain largely undescribed.

*Caenorhabditis elegans* is a powerful model system for investigating axon regeneration and developmental axon guidance^16–27^. The animal’s relatively small, fully mapped nervous system and transparent cuticle enable axotomy of individual, defined axons, followed by in vivo visualization of axon regeneration and guidance. The *C. elegans* genome is highly conserved with mammals, and several key regulators of axon regeneration, including DLK-1/DLK and DAF-18/PTEN, function similarly across species^5,16,17,19,28,29^. The tractability of *C. elegans* also enabled the seminal discovery of conserved developmental guidance cues such as UNC-6/Netrin and its receptors UNC-40/DCC and UNC-5/UNC5C^22–25^. Because wild-type *C. elegans* motor axons are capable of regenerating and are directed towards post-synaptic muscles along the dorsal nerve cord, we reasoned that this system would be an informative model to investigate how regenerating axons navigate the mature nervous system.

Here, we identify a post-developmental guidance mechanism that directs regenerating motor axons alongside intact neuronal processes. We find that injured cholinergic axons, and not GABA axons, preferentially regenerate adjacent to the branched dendrites of the PVD mechanosensory neuron. This form of guidance appears specific to the mature nervous system; during development, cholinergic axons extend and form synapses with their postsynaptic partners before the PVD neuron elaborates its dendrites^30–32^. We find that Nidogen (NID-1), a conserved basement membrane protein, helps guide regenerating cholinergic axons alongside PVD dendrites toward their pre-injury locations and promotes functional recovery. NID-1 interacts with integrin and laminin to guide cholinergic axons in a cell-type-specific manner: regenerating GABA axons do not normally extend alongside the PVD dendrites, but do so when integrin is ectopically expressed. These findings reveal a mechanism of post-developmental guidance that can be used to redirect axons in the mature nervous system.

## Results

### Regenerating cholinergic motor axons colocalize with PVD dendrites

During *C. elegans* development, cholinergic and GABAergic motor axons extend circumferentially from their cell bodies in the ventral nerve cord to the dorsal nerve cord, adopting a relatively straight morphology that is maintained through adulthood (Figure 1A). After laser axotomy, some motor axons regenerate directly back to their original target; however, many regenerate in a less direct path and make sharp 90-degree turns (Figure 1B,C). We observed that the morphology of these regenerating axons resembles the highly branched dendrites of a multimodal sensory neuron called the PVD (Figure 1D,E,F).

**Figure 1.**
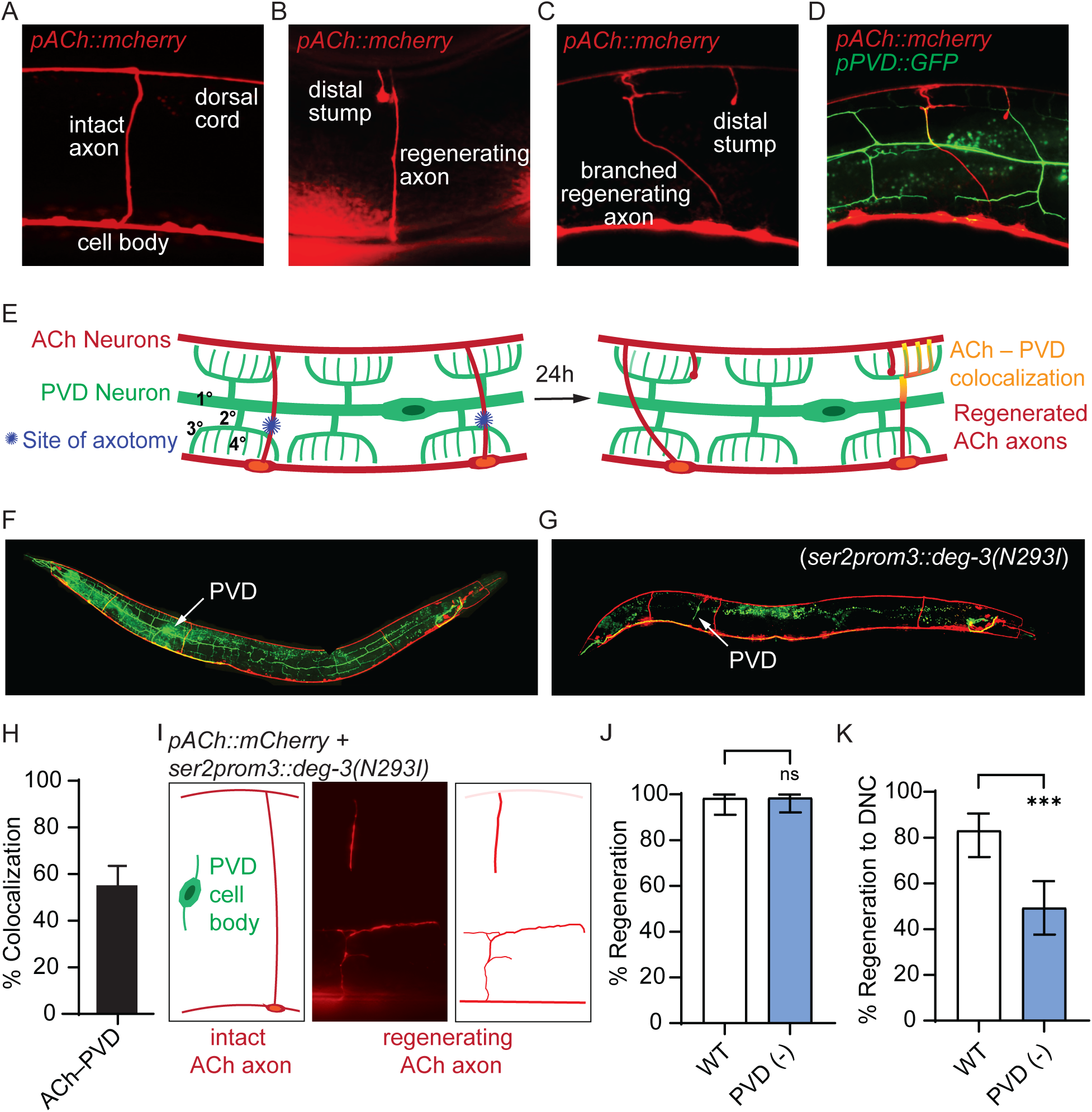
PVD dendrites increase the ability of regenerating cholinergic axons to reach the dorsal nerve cord. **(A)** Wild-type mCherry-labeled cholinergic motor axon (*Pacr-2::mCherry*) extending circumferentially from its cell body in the ventral nerve cord to the dorsal nerve cord of *C. elegans*. **(B)** Regenerated cholinergic motor axon with morphology similar to uninjured wild-type axons. The distal stump is the portion of the axon severed from the cell body upon injury and confirms that the axon was successfully severed. **(C)** Regenerated cholinergic axon with a branched morphology. **(D)** Regenerated cholinergic axon colocalized with GFP-labeled PVD dendrites (*F49H12.4p::GFP*). **(E)** Axotomy model depicting primary, secondary, tertiary and quaternary dendritic branches of the PVD (green), ACh axons (red), site of axotomy (blue; always ventral to the primary branch of the PVD), and colocalization between a regenerating ACh axon and PVD dendrites (yellow). **(F)** Otherwise wild-type animal expressing *Pacr-2::mCherry* (ACh neurons, red) and *F49H12.4p::GFP* (PVD neurons, green). **(G)** Expression of *ser-2prom3::deg-3-N293I*) genetically ablates PVD dendrites. **(H)** 55.12% of regenerating ACh axons colocalize with PVD dendrites (n=127). **(I)** Representative cartoon of uninjured cholinergic axon and a PVD neuron that does not extend dendrites. Representative micrograph and tracing of a regenerating ACh axon that fails to reach the dorsal cord in the absence of PVD dendrites. **(J)** Genetic PVD ablation (*ser-2prom3::deg-3-N293I*) does not alter the proportion of injured axons that initiate regeneration relative to wild type (N=60, 68). **(K)** Despite having similar numbers of regenerating ACh axons, significantly fewer reach the dorsal cord when PVD dendrites are absent compared to wild-type animals (N=59, 67). Significance determined by Fisher’s exact test; *p≤0.05, **p≤0.01, and ***p≤0.001. Error bars represent 95% confidence intervals.

To determine whether, in the mature nervous system, regenerating ACh axons colocalize with PVD dendrites, individual mCherry-labeled cholinergic motor axons were axotomized slightly below the lateral midline, without damaging GFP-labeled PVD dendrites (Figure 1E). All axotomies were performed in late L4 animals, a stage at which cholinergic and GABAergic motor neurons have completed their dorsal migrations and PVD dendrites have elaborated their arbor. Regeneration was assessed 24 hours later in young adults. Thus, regeneration phenotypes described here occur in a mature environment rather than during developmental circuit formation. We found that twenty-four hours after injury, 55% of regenerating cholinergic motor axons were colocalized with PVD dendrites (Figure 1H). Here, we define colocalization as overlap of fluorescent signal between the regenerating mCherry-labeled ACh axon and GFP-labeled PVD dendrites over at least 2.5 μm, equivalent to 5% of the width of adult *C. elegans*. This criterion excludes axons that cross but do not follow PVD dendrites. We observed colocalization between regenerating ACh axons and multiple levels of dendritic branches and once a regenerating axon colocalized with the PVD at any branch level, it frequently continued to colocalize with subsequent levels of dendritic branches, indicating regenerating ACh axons are not restricted to a specific type of dendritic branch or associated extracellular environment (Figure S1A). These data suggest the presence of PVD dendrites may help guide regenerating cholinergic motor axons to the dorsal nerve cord in the mature nervous system.

### The presence of PVD dendrites increases the ability of regenerating cholinergic axons to reach the dorsal nerve cord

We asked whether PVD dendrites are required for cholinergic axon regeneration by repeating the axotomy experiment in animals expressing a tissue-specific degron that genetically ablates the PVD dendrites (*ser-2prom3*::*deg-3-N293I*), hereafter referred to as PVD-ablated^33^ (Figure 1G, I-K). The absence of PVD dendrites did not alter the proportion of cholinergic axons that initiate regeneration compared with wild-type animals (Figure 1J); however, significantly fewer regenerating cholinergic axons reached the dorsal nerve cord (Figure 1K). These data indicate that the presence of PVD dendrites provides physical or molecular cues that direct regenerating cholinergic axons to the dorsal nerve cord.

Interestingly, a substantial proportion of cholinergic axons still reached the dorsal nerve cord in PVD-ablated animals (Figure 1K), suggesting additional cues may orient regenerating axons. To determine which trajectories are used in the absence of PVD dendrites, we axotomized mCherry-labeled cholinergic axons in animals expressing P*rgef-1*::GFP, a pan-neuronal GFP reporter (Figure 2A). We found that in the presence of PVD dendrites, only 11.54% of regenerating cholinergic axons colocalized with uninjured GABA commissures (Figure 2B), indicating that regenerating ACh axons do not preferentially follow the trajectories of GABAergic commissures when PVD dendrites are available. However, in PVD-ablated animals, significantly more regenerating cholinergic axons colocalized with intact GABA commissures (Figure 2B), indicating regenerating ACh axons can extend alongside GABAergic commissures in the absence of PVD dendrites, yet the trajectory along PVD dendrites is preferred. The majority of ACh axons that colocalized with the GABA axons successfully reached the dorsal cord (Figure 2C), demonstrating that regenerating ACh axons can reach the dorsal nerve cord along GABAergic trajectories. Together, these findings indicate cholinergic motor axons can be guided back to the dorsal nerve cord along or in the immediate vicinity of uninjured neuronal processes and preferentially follow trajectories associated with PVD dendrites rather than GABAergic commissures.

**Figure 2.**
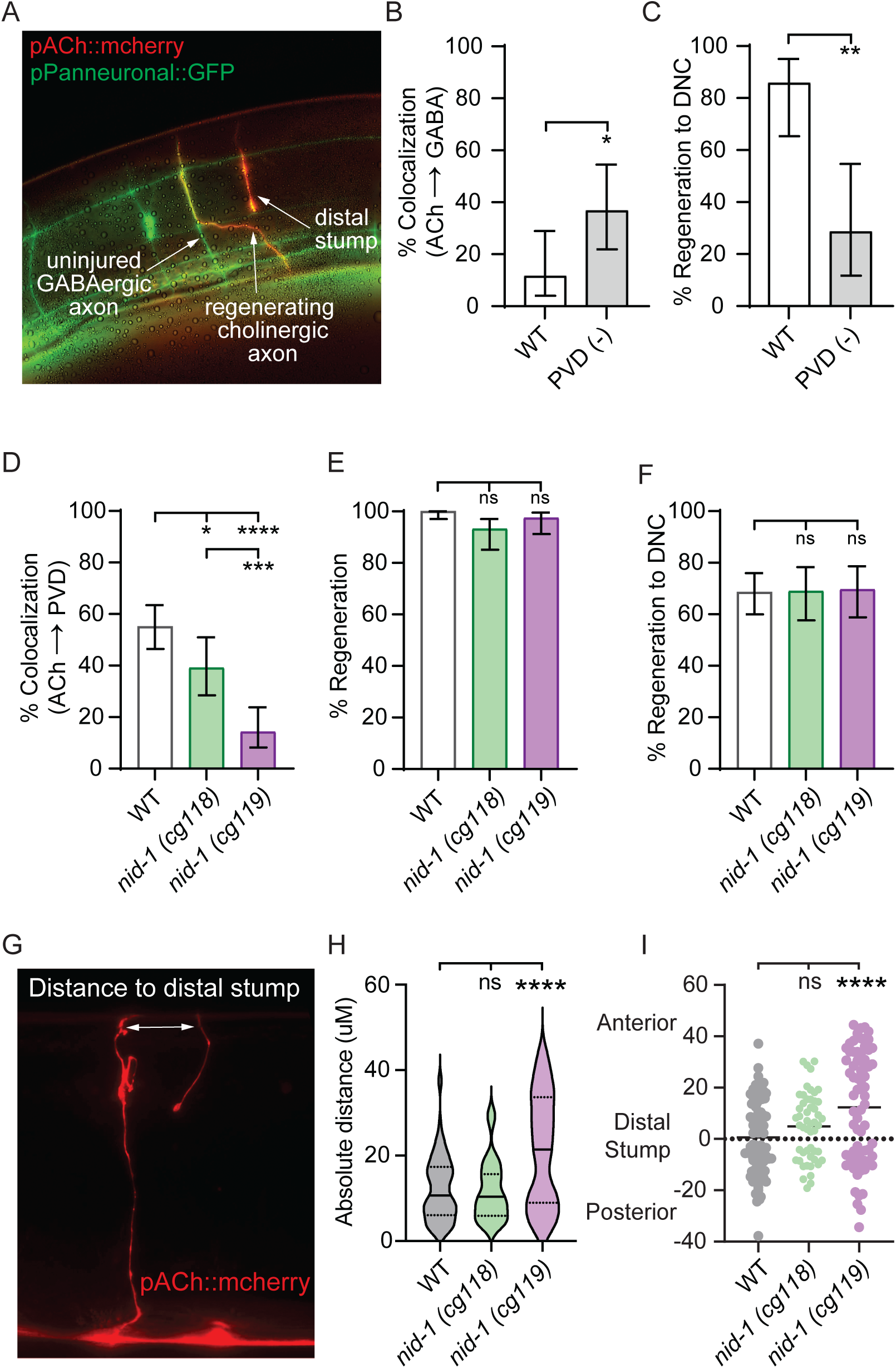
Regenerating cholinergic motor axons colocalize with PVD dendrites, which increase their ability to reach the dorsal nerve cord. **(A)** Representative image of an injured regenerated ACh axon colocalized with a GABAergic motor axon in the absence of PVD dendrites. Animals express *Pacr-2::*mCherry to label ACh neurons and the pan-neuronal GFP marker, *Prgef-1::*GFP to label all other neurons. GABA neurons are readily identified in this strain as GFP-labeled ventral-dorsal directed commissures. **(B)** The number of regenerating ACh axons that colocalize with GABA axons is increased in the absence of PVD dendrites (N=26,30). **(C)** In wild-type animals, the majority of regenerating axons that colocalize with GABA commissures reach the dorsal nerve cord. (N=21,14). **(D)** Fewer regenerating ACh axons colocalize with PVD dendrites in *nid-1(cg118)* and *nid-1(cg119)* mutants. The null *cg119* mutation disrupts the ACh-PVD interaction more significantly than the hypormophic *cg118* mutation (N=127,69,77). **(E, F)** Neither mutation disrupts the proportion of injured axons that initiate regeneration, the proportion of regenerating axons that reach the dorsal cord, or the proportion of PVD dendrites that colocalize with intact ACh axons during development (N=127,74,79). **(G)** Representative image of the measurement of distance to distal stump **(H)** Among the injured regenerated ACh axons that reach the dorsal cord, the distance between where the regenerated axons join the dorsal cord and their respective distal stump (distance to distal stump) is greater in animals lacking functional NID-1 (*nid-1(cg119)*) (N=84,73,51). **(I)** In animals lacking NID-1 (*nid-1(cg119)*), the insertion point of regenerated ACh axons into the dorsal cord is anteriorly biased (N=84,73,51). Significance determined by Fisher exact test for (B-G), Mann-Whitney for (H, I) and is indicated by *≤0.05, **p≤0.01, ***p≤0.001, and ****p≤0.0001. Error bars represent 95% confidence interval.

### NID-1/Nidogen guides regenerating cholinergic axons alongside the PVD

The directed pattern of cholinergic motor axon regeneration suggests that molecular cues associated with PVD dendrites, or their surrounding extracellular environment, guide regenerating axons. We asked whether conserved basement membrane proteins might be repurposed after development to guide regenerating ACh axons. We found that loss of function mutations in *nid-1*, a highly conserved basement membrane glycoprotein homologous to Nidogen in mammals, significantly reduced the number of regenerating axons that colocalized with PVD dendrites compared to wild-type regenerating axons (Figure 2D). Colocalization was more strongly disrupted in animals carrying the *cg119* deletion allele, which is a molecular null, compared with animals carrying the hypomorphic *cg118* allele, an in-frame deletion that disrupts two of three globular domains that bind basement membrane proteins. Notably, the proportion of injured axons that regenerated was not altered by loss of *nid-1* function, indicating the disrupted colocalization was not caused by an inability to regenerate (Figure 2E). In uninjured animals, loss of *nid-1* function did not grossly alter the morphology of ACh commissures (Figure S2A), nor did it alter the frequency or pattern of interaction between developing PVD dendrites and established motor axons (Figure S2B,C), which is restricted to secondary dendrites^30^. Together, these data indicate that NID-1 helps guide regenerating cholinergic axons along PVD-like trajectories in the mature nervous system and does not determine their ability to regenerate.

### NID-1 promotes synapse reformation after injury

During development, motor axons extend circumferentially from the ventral nerve cord to the dorsal nerve cord, where they turn longitudinally and form en passant synapses with body wall muscles. After injury, the location at which each axon originally entered the dorsal nerve cord can be determined by visualizing the severed distal stump, which does not degenerate in *C. elegans* and therefore provides a visible landmark of the axon’s pre-injury position. We found that although ACh axons regenerate to the dorsal nerve cord in *nid-1(-)* animals (Figure 2F), they contact the dorsal nerve cord significantly further away from their severed distal stump in *nid-1(cg119)* null mutants compared with wild-type animals (Figure 2G,H). A more detailed analysis revealed that whereas wild-type axons reach the dorsal cord on either side of the distal stump, there is an anterior bias in *nid-1(cg119)* null mutants: regenerated axons are more likely to meet the dorsal cord anterior to the distal stump (Figure 2I). In contrast, the distance between regenerating *nid-1(cg118)* axons and the distal stump was not significantly different, consistent with the milder PVD-colocalization phenotype of this hypomorphic allele (Figure 2D). These data indicate that loss of *nid-1* alters the location at which regenerating ACh axons return to the dorsal nerve cord. Since the functional significance of such a positional shift is not known, the consistent change in trajectory motivated us to investigate whether loss of *nid-1* function also affects the axon’s ability to reform functional synapses with the appropriate cells.

To assess whether regenerated ACh axons re-establish synapses along the dorsal nerve cord, we examined the distribution of synaptic vesicle marker RAB-3, a widely used reporter of presynaptic sites^34^. The photoconvertible P*unc-129*::Dendra2::RAB-3 reporter, which is expressed specifically in cholinergic neurons, was used to distinguish newly formed synaptic vesicle puncta^35^. Before photoconversion, green Dendra2::RAB-3 puncta cluster along cholinergic axons in the dorsal nerve cord where cholinergic axons form *en passant* synapses with GABAergic neurons and body wall muscles (Figure 3A). When photoconverted with 405nm light, existing Dendra2::RAB-3 proteins emit red fluorescence, distinguishing them from newly formed green fluorescent synaptic vesicles. To quantify newly formed synapses, we severed individual cholinergic axons and then irreversibly photoconverted Dendra2::RAB-3. Twenty-four hours after injury and photoconversion, several regenerating axons had reached the dorsal cord. At this timepoint, newly formed RAB-3 puncta should appear as green fluorescence in the portion of regenerated axon that extended along the dorsal cord. Prior to injury, we observed a similar number of RAB-3 puncta along the dorsal cords of *nid-1(cg119)* null and wild-type animals (Figure 3B). However, regenerated *nid-1(cg119)* axons contained significantly fewer RAB-3 puncta along the dorsal nerve cord compared to wild-type animals (Figure 3C), indicating NID-1 is important for synapse reformation. While exposure to 405nm light during photoconversion did not affect the proportion of axons that initiated regeneration (Figure S3A), it significantly decreased the proportion of regenerating axons that reached the dorsal cord 24 hours after axotomy (Figure S3B). We hypothesize the 405nm wavelength light induces the production of reactive oxygen species and oxidative stress^36–38^. Nonetheless, the difference in number of RAB-3 puncta between *nid-1* null and wild-type animals reflects the requirement for *nid-1* function, as the photoconversion dynamics were held constant between *nid-1(-)* animals and wild-type controls.

**Figure 3.**
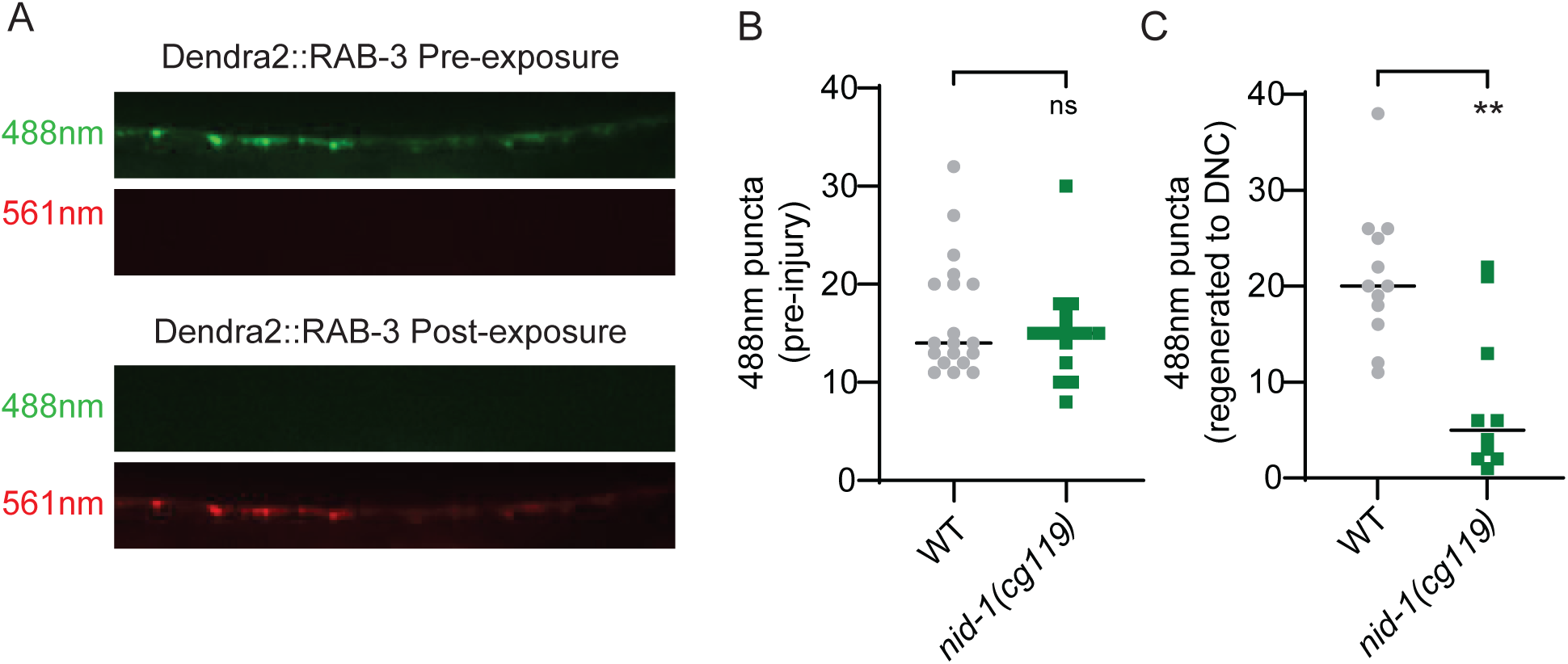
NID-1 promotes synapse formation after injury. **(A)** Representative image of RAB-3 tagged with Dendra2 before (green) and after photoconversion (red). **(B)** *nid-1(cg119)* mutants and wildtype animals have a similar number of RAB-3 puncta before injury (N=19,16). **(C)** In *nid-1(cg119)* animals, regenerating axons that reach the dorsal nerve cord regain significantly fewer synaptic RAB-3 puncta compared to wildtype animals, 24 hours after injury (N= 12,10). Significance determined by Mann-Whitney, **p≤0.01.

### Expressing NID-1 in the body wall muscle, not the hypodermis, is sufficient to rescue synapse reformation

NID-1 is a diffusible basement membrane glycoprotein expressed primarily in the hypodermis and body wall muscle and distributed throughout the animal^39^. To determine where NID-1 functions to guide regenerating ACh axons, we conducted tissue-specific rescue experiments. Specifically, we expressed the NID-1A isoform driven by either cholinergic neuron-, body wall muscle-, or hypodermis-specific promoters (*Punc-17, Pmyo-3, Pdpy-7,* respectively) in animals that otherwise lack *nid-1* function (*nid-1(cg119)),* and quantified axon guidance. We performed these experiments with the NID-1A isoform because it is the longest of the three endogenous NID-1 isoforms in *C. elegans*, contains all predicted functional domains, and is most conserved with mammalian nidogens^39^. The two shorter NID-1B and NID-1C isoforms differ only in alternative splicing events that affect the rod domain, which is a flexible linker between N- and C-terminal domains^39–42^.

We found that cholinergic motor neuron-specific NID-1A expression (*Punc-17::nid-1A*) did not rescue colocalization between regenerating ACh axons and the PVD (Figure 4A). In contrast, tissue-specific expression of NID-1A in either the hypodermis or body wall muscle of *nid-1(cg119)* animals was sufficient to restore colocalization with PVD dendrites (*Pmyo-3::nid-1A* and *Pdpy-7::nid-1A*, respectively). At higher resolution, tissue-specific expression of NID-1A in either the body wall muscle or the hypodermis, but not ACh neurons, guided regenerating ACh axons to their distal stump (Figure 4B). None of the NID-1A transgenes influenced the proportion of injured axons that initiated regeneration, including ectopic expression in the cholinergic motor neurons (Figure 4C), suggesting NID-1 regulates axon guidance rather than the ability to regenerate. Together, these data indicate that NID-1A functions cell non-autonomously to guide regenerating ACh axons alongside the trajectory of PVD dendrites and reach their pre-injury locations.

**Figure 4.**
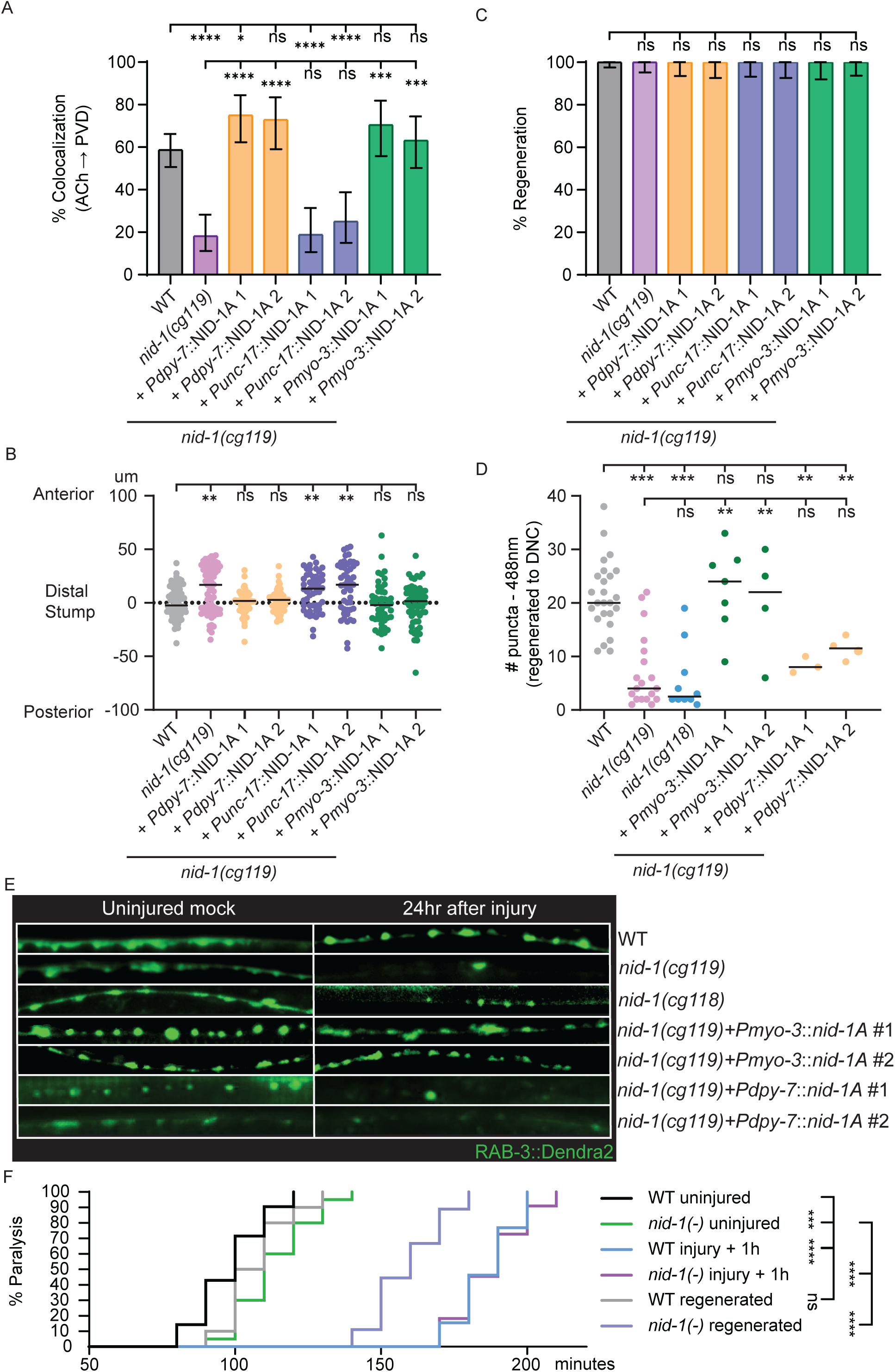
Tissue-specific NID-1 regulates regenerating axon guidance and synapse formation. **(A)** Expressing NID-1A in the hypodermis or body wall muscles rescues colocalization between regenerating ACh axons and PVD dendrites in *nid-1(cg119)* animals. Ectopic expression of NID-1A in the ACh neurons does not rescue the phenotype (N = 150, 77, 56, 48, 53, 48, 44, 57). **(B)** Expressing NID-1A in body wall muscles and the hypodermis rescues the distance between injured regenerated ACh axons and their distal stumps. Ectopic expression of NID-1A in ACh neurons does not rescue guidance (N=84,73, 51,54,48,53,48,48,57). **(C)** Expressing NID-1A in the hypodermis or body wall muscles does not affect the proportion of axons that regenerate (N = 150, 77, 56, 48, 53, 48, 44, 57). **(D)** Fewer RAB-3 puncta are formed in injured regenerated ACh axons in *nid-1(cg119)* and *nid-1(cg118)* mutants relative to wild-type. Expressing NID-1A in body wall muscles rescues the number of RAB-3 puncta (N=25,19,10,7,4,3,4). **(E)** Green Dendra2::RAB-3 puncta formed in regenerated axons after existing puncta have been photoconverted to red. Micrographs represent puncta in mock injured and regenerated ACh axons in wildtype, *nid-1(cg119)*, *nid-1(cg118)*, body wall-specific NID-1A rescue, and hypodermis specific NID-1A rescue animals. **(F)** Nine commissural cholinergic axons were injured in each animal, 3 on the right side of the animal and 6 on the left side. Uninjured *nid-1(cg119)* null animals (green) become paralyzed on aldicarb slightly after wildtype animals (black), reflecting a mild developmental synaptic defect in *nid-1* animals. One hour after injury, both WT (blue) and *nid-1(-)* animals are severely resistant to aldicarb, indicating cholinergic function has been disrupted by axotomy. 24 hrs post-injury, WT animals (grey) recover pre-injury (black) levels of susceptibility to aldicarb. However, *nid-1(-)* animals fail to recover pre-injury levels of function (green). (N=21,20,13,11,10,9). Significance determined by Fisher exact test (A,B), Kruskal-Wallis (C), Mann-Whitney (D) and Mantel–Cox (log-rank) test (F), and is indicated by *≤0.05, **p≤0.01, ***p≤0.001, and ****p≤0.0001. Error bars represent 95% confidence intervals.

To determine where NID-1A expression is sufficient to restore synapse reformation, we crossed Dendra2::RAB-3 into each of the NID-1A-expressing strains that restored wild-type guidance. As in the previous Dendra experiments, we photoconverted Dendra and quantified the number of newly formed green RAB-3 puncta 24 hours after axotomy. Body wall muscle-specific expression of NID-1A restored the number of Dendra2::RAB-3 puncta to regenerated ACh axons along the dorsal nerve cord, whereas hypodermis-specific-NID-1A expression did not (Figure 4D, 4E). Therefore, muscle-derived NID-1A is sufficient to support synapse reformation after injury. Although hypodermis-specific expression was sufficient to rescue axon guidance, it did not rescue Dendra2::RAB-3 puncta reformation (Figure 3D, 3E). These data suggest synapse reformation is regulated by muscle-derived NID-1A.

### NID-1 promotes functional axon regeneration

Our finding that NID-1 promotes synaptic RAB-3 puncta reformation after injury suggests NID-1 is also required for functional recovery. To test this hypothesis, we performed aldicarb assays on *nid-1(cg119)* and wild-type animals before and after injury. Aldicarb is an acetylcholinesterase inhibitor that causes acetylcholine to accumulate at functional neuromuscular junctions, leading to paralysis. When exposed to aldicarb, uninjured *nid-1(-)* animals became paralyzed more slowly than wild-type animals (Figure 4F), agreeing with previously published data that *nid-1(-)* mutants have developmental defects in synapse function^43^. We axotomized nine commissural cholinergic axons, six on the left side and three on the right side of each animal. Animals were first transferred to standard NGM plates to recover, then transferred to 1 mM aldicarb plates 1 hour or 24 hours after injury. At 1 hour, animals from both genotypes were more resistant to aldicarb than uninjured controls, confirming that axotomy successfully disrupted cholinergic motor function (Figure 4F). Wild-type animals that had recovered on NGM plates for 24 hours paralyzed at the same rate as uninjured wild-type animals, indicating they regained ACh function. However, *nid-1(-)* animals remained significantly resistant to aldicarb, paralyzing more slowly than injured wild-type animals and uninjured *nid-1(-)* mutants. These data indicate *nid-1* is required for functional repair and that the functional consequence of losing *nid-1* function is more severe during regeneration than during development. Together with the observation that *nid-1(-)* and wild-type animals regenerate the same proportion of injured axons to the neuromuscular junction (Figure 2F), and that *nid-1*(-) axons regain fewer RAB-3 puncta (Figure 3C), these findings indicate functional synapse reformation is disrupted in *nid-1(-)* animals.

### Integrin guides regenerating cholinergic axons

We next asked whether regenerating ACh motor neurons are preferentially guided along PVD dendrites compared to GABA motor neurons. Like ACh axons, regenerating GABA axons also extend circumferentially from the ventral to the dorsal nerve cord, and are localized as near to PVD dendrites as ACh axons; however, we found that regenerating GABA axons colocalized with PVD dendrites significantly less frequently than ACh axons (Figure 5A). These data suggest the mechanisms that mediate colocalization must differ, at least in part, between cholinergic and GABAergic neurons.

**Figure 5.**
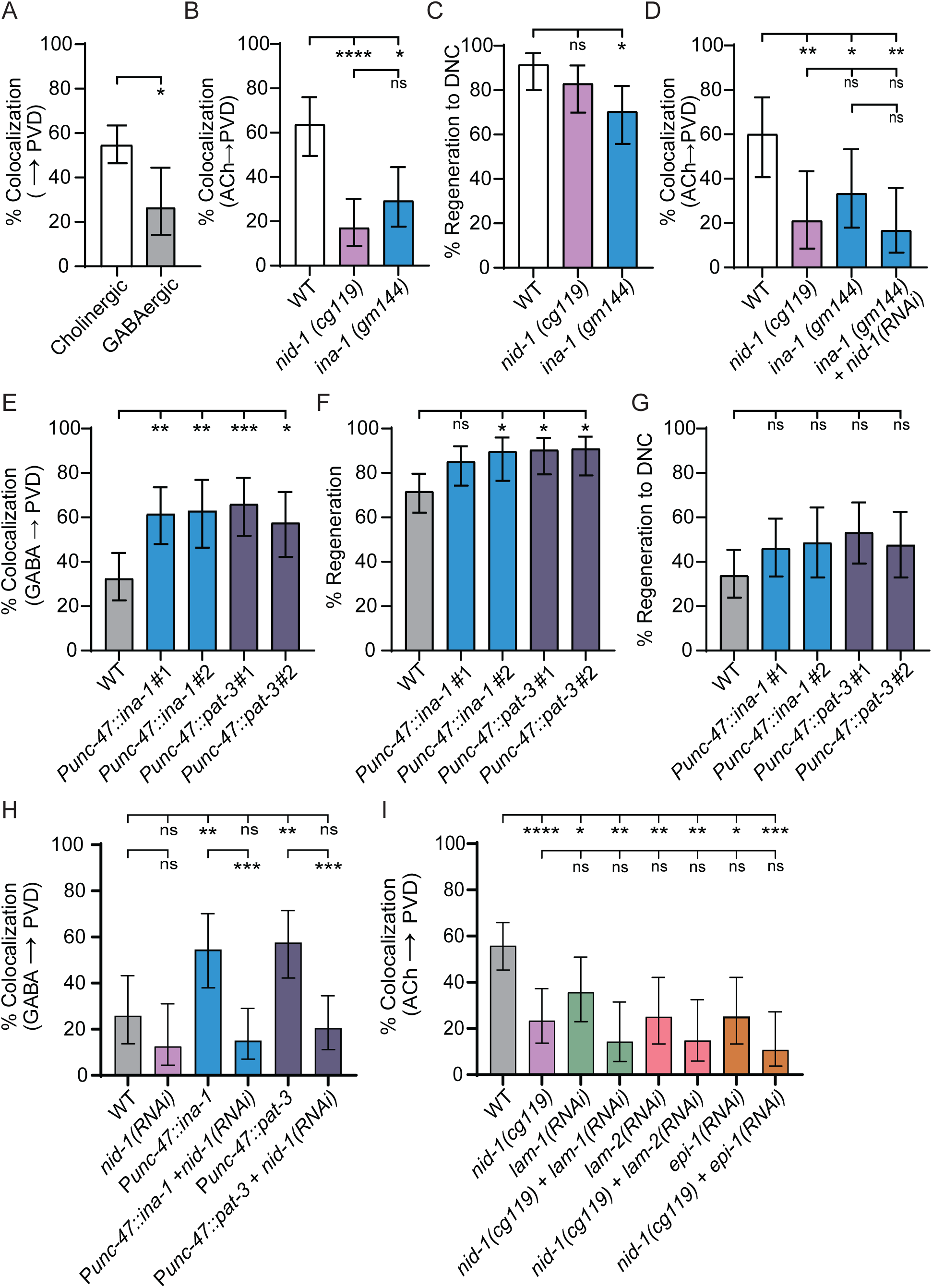
Integrin and laminin promote guidance of regenerating ACh axons. **(A)** Regenerating GABA axons colocalize with PVD dendrites significantly less frequently compared to regenerating ACh axons (N=127,30). **(B)** The *ina-1 (gm144)* mutation disrupts colocalization of regenerating ACh axons and PVD dendrites compared to wildtype animals. The frequency of colocalization is similar to that of *nid-1(cg119)* mutants (N=47,47,41). **(C)** Regenerating ACh axons reach the dorsal cord less frequently in *ina-1(gm144)* mutant animals compared to wild type (N=47, 47, 44). **(D)** Feeding *nid-1* RNAi to wild-type animals decreases colocalization between regenerating ACh axons and PVD dendrites. However, it does not significantly enhance the *ina-1(gm144)* mutant phenotype (N=25, 19, 24, 24). **(E)** Overexpressing integrin subunits INA-1 and PAT-3 in GABAergic axons significantly increases colocalization between regenerated GABAergic axons and PVD dendrites compared to wild-type (N=71, 52, 35, 47, 40). **(F)** Overexpressing integrin subunits INA-1 and PAT-3 in GABAergic axons significantly increase the proportion of GABAergic axons that initiate regeneration compared to wild-type (N=99,61,39,52,44). **(G)** Overexpressing integrin subunits INA-1 and PAT-3 in GABAergic axons does not increase the proportion of regenerating GABAergic axons that reach the dorsal cord (N=71,52,35,47,40). Significance determined by Fisher exact test and is indicated by *≤0.05, **p≤0.01, ***p≤0.001, and ****p≤0.0001. Error bars represent 95% confidence interval. **(H)** NID-1 RNAi suppresses colocalization between regenerating GABAergic axons and PVD dendrites in animals that over express integrin subunits INA-1 and PAT-3 (N=31,24,33,40,40,44). **(I)** RNAi targeting *lam-1*, *lam-2*, or *epi-1* decreases the frequency with which regenerating ACh axons colocalize with PVD dendrites (N=86,47,42,28,32,27,32,28). Significance determined by Fisher exact test and is indicated by *≤0.05, **p≤0.01, ***p≤0.001, and ****p≤0.0001. Error bars represent 95% confidence interval.

To identify ACh-specific proteins that might mediate guidance, we analyzed the expression of candidate cell-surface proteins across GABA, ACh, and PVD neurons using the CENGen (*C. elegans* Neuronal Gene Expression Map & Network) single-cell RNA-sequencing database^44^. The comparison revealed that ACh neurons express significantly higher levels of integrin compared to GABA neurons^44,45^. *C. elegans* express three integrin subunits, INA-1, PAT-2, and PAT-3, which heterodimerize to form two integrin complexes: α INA-1/β PAT-3 and α PAT-2/β PAT-3. These complexes regulate several functions, including axon outgrowth and synapse maintenance during development, by forming cell-surface receptors that communicate signals from the extracellular matrix to intracellular signaling pathways^46^.

The difference in integrin expression between ACh and GABA neurons prompted us to ask whether integrin contributes to guiding regenerating axons. A loss-of-function mutation in the integrin α subunit *ina-1* significantly reduced the proportion of regenerating ACh axons that colocalized with PVD dendrites (Figure 5B). This phenotype was similar to that observed in *nid-1* mutants, suggesting integrin and nidogen may act in a shared pathway to guide regenerating axons in the mature nervous system. We were unable to directly test whether *nid-1* and *ina-1* function in the same pathway by generating *nid-1(cg119)*; *ina-1(gm144)* double mutants, because the combination is lethal. Alternatively, we knocked down *nid-1* function in *ina-1(gm144)* mutants using *nid-1* RNAi. We did not observe any further reduction in the number of regenerating ACh axons that colocalize with PVD dendrites compared to either single mutant (Figure 5D). This negative result is not due to a lack of RNAi efficiency, since *nid-1* RNAi in an otherwise wild-type background disrupted colocalization to the same extent as the genetic null mutation *cg119*. Therefore, our data suggest *nid-1* and *ina-1* function in the same genetic pathway to guide regenerating ACh axons. Neither *nid-1* nor *ina-1* loss of function affected regeneration initiation (Figure S5A,B), though *ina-1(gm144)* mutants showed a modest reduction in the proportion of axons reaching the dorsal cord (Figure 5C), indicating that the colocalization defect reflects a guidance phenotype rather than a general inability to regenerate.

### Ectopic integrin expression is sufficient to guide regenerating GABA motor axons

We asked if integrin expression levels determine whether regenerating motor axons colocalize with PVD dendrites. To do so, we overexpressed the integrin subunits *ina-1* (P*unc-47*::*ina-1)* and *pat-3* (P*unc-47*::*pat-3)* in GABA neurons and quantified colocalization between regenerating GABA axons and PVD dendrites. Twenty-four hours after axotomy, we observed that cell-autonomous overexpression of either PAT-3 or INA-1 increased colocalization between regenerating GABAergic axons and PVD dendrites (Figure 5E). Integrin expression also increased the proportion of injured GABA axons that regenerate (Figure 5F), but did not increase the proportion of regenerating axons that extend back to the dorsal cord (Figure 5G). Together, these data indicate that integrin expression determines cell-type-specific guidance of regenerating axons in the mature nervous system, and that this mechanism can be co-opted to redirect regenerating axons in the mature nervous system.

### Integrin-induced guidance requires NID-1

Since we discovered that integrin overexpression can increase colocalization between regenerating axons and PVD dendrites, we hypothesized that integrin requires NID-1 to mediate this interaction. We found that feeding *nid-1* RNAi to animals that overexpress INA-1 or PAT-3 in their GABA axons significantly decreased the proportion of regenerating GABAergic motor axons that colocalized with PVD dendrites (Figure 5H). However, the frequency of regeneration was not affected by RNAi (Figure S6A, S6B). These data indicate that the NID-1-integrin interaction is required to guide regenerating motor axons in mature animals.

### Laminin promotes guidance of regenerating cholinergic axons

Laminin is a major component of the basement membrane and a known binding partner of Nidogen^47^. It scaffolds various cell surface receptors and regulates basement membrane integrity and extracellular matrix organization during development. The *C. elegans* genome encodes four laminin subunits (α: EPI-1 and LAM-3; β: LAM-1; and γ: LAM-2). Mutations in these structural components lead to misassembled and/or broken basement membranes, resulting in severe defects in tissue organization, temperature sensitivity, and embryonic or larval lethality^48–50^. As strong loss-of-function mutations in laminin subunits cause severe developmental defects including embryonic or larval lethality^51^, we knocked down the laminin subunits *lam-1*, *lam-2*, and *epi-1* using RNAi. We found that loss of *lam-1*, *lam-2*, or *epi-1* function significantly decreased the proportion of regenerating cholinergic axons that colocalized with PVD dendrites compared to wild-type controls. However, the phenotypes were not significantly different from those observed in *nid-1(cg119)* null animals (Figure 5I). Loss of *lam-2* function was the only laminin subunit that caused a significant decrease in the proportion of regenerating axons that reach the dorsal cord (Figure S6C, S6D). Taken together, our data reveal a tripartite nidogen-laminin-integrin pathway that guides regenerating cholinergic axons in the mature nervous system. These results suggest that manipulating expression of this complex can redirect other cell-types along defined paths.

## Discussion

A central challenge in repairing the injured nervous system is understanding how surviving axons navigate a complex post-developmental environment to reach their targets. Although some developmental guidance cues persist into adulthood, relatively little is known about the mechanisms that direct regenerating axons after development. Here, we identify a post-developmental guidance mechanism in which regenerating cholinergic axons extend alongside intact neuronal processes to reach their pre-injury location in the dorsal nerve cord. This process requires the basement membrane protein NID-1/Nidogen, together with laminin and integrin, to guide axons along appropriate trajectories.

### Regenerating axons extend alongside PVD dendrites in the mature nervous system

Our results indicate that regenerating axons preferentially extend alongside intact neuronal processes. In wild-type animals, regenerating cholinergic motor axons frequently align with the dendritic arbor of the PVD sensory neuron as they grow toward the dorsal nerve cord (Figure 1H). When PVD dendrites are absent, regenerating axons instead extend alongside intact GABAergic commissures (Figures 1K, 2A–C). These observations suggest that intact neuronal architecture influences the direction of axon regeneration in the mature nervous system and that molecular cues preferentially guide axons alongside PVD dendrites. Since cholinergic axons develop before PVD dendrites are elaborated^30–32^, this pattern of colocalization suggests that the mechanism responsible for guiding regenerating axons must differ, at least in part, from that used during development.

### NID-1/Nidogen mediates guidance of regenerating axons alongside intact neuronal processes

Nidogen is a diffusible basement membrane protein that contributes to extracellular matrix organization across species^52–54^. In *C. elegans*, NID-1 is expressed primarily by non-neuronal tissues adjacent to motor neurons, including the body wall muscle and hypodermis^39,44^. Loss of *nid-1* function significantly reduced colocalization between regenerating ACh axons and PVD dendrites (Figure 2D), and caused regenerated axons to contact the dorsal nerve cord further from their pre-injury position (Figures 2G,H). Our tissue-specific rescue experiments further indicate that expression of NID-1 in surrounding non-neuronal tissues is sufficient to restore accurate axon guidance (Figures 4A,C), consistent with NID-1 acting as a diffusible extracellular cue in the local microenvironment of regenerating axons. A key feature of this guidance mechanism is that NID-1’s role in directing cholinergic axons alongside PVD dendrites is specific to regeneration. In uninjured animals, *nid-1* mutant motor axons develop normally: their initial dorsoventral pathfinding is unaffected (Figure S2A), and the frequency and spatial pattern of colocalization between motor axons and PVD dendrites are indistinguishable from wild type (Figure S2B,C). NID-1 is secreted by the body wall muscle and hypodermis^39^ and has established roles in positioning longitudinal axon tracts^55^ and promoting PVD secondary dendrite outgrowth during larval development^56^. Yet regenerating axons preferentially associate with higher-order branches of the PVD arbor (Figure S1B), in addition to the secondary branches^56^, suggesting that the injury response engages NID-1 in a distinct spatial context. Whether regenerating axons respond to NID-1 through direct contact with PVD dendrites or through diffuse ECM cues distributed along their trajectories remains an open and important question.

### Nidogen promotes synapse reformation and functional axon regeneration

Functional axon regeneration requires not only axon growth and guidance but also the re-establishment of synaptic connections. We found that loss of *nid-1* function impairs recovery of presynaptic vesicle puncta along the dorsal nerve cord (Figure 3C) and functional recovery of regenerating cholinergic axons as assessed by aldicarb assay (Figure 3F), indicating that NID-1 contributes to multiple aspects of functional axon regeneration. NID-1 contains three globular domains called G1, G2, and G3, which are joined by a rod domain connecting G1/G2 to G3. Comparison of phenotypes between the *nid-1(cg118)* hypomorphic allele, which disrupts the G1 and G2 domains while leaving G3 intact, and the *nid-1(cg119)* null allele, reveals that axon guidance and synapse reformation have distinct requirements for *nid-1*. The *nid-1(cg118)* mutants have only mild axon guidance defects compared to null mutants (Figures 2D, S2A), yet synapse reformation is similarly impaired in both alleles (Figures 3C, 4D,E), suggesting that synapse reformation is more sensitive to loss of NID-1 function than axon guidance. Whether this reflects a difference in sensitivity to levels of NID-1 or a specific requirement for the G1 and G2 domains in synapse reformation remains to be determined. The spatial requirements for NID-1 also differ between the two functions. Expression of NID-1 in either body wall muscles or the hypodermis is sufficient to rescue axon guidance defects (Figures 4A,C), whereas synapse reformation requires NID-1 expression specifically in body wall muscles (Figures 4D,E). NID-1 may therefore function in two capacities: as a diffusible guidance cue during axon extension, and, consistent with evidence that Nidogen anchors synapses to the basement membrane during development^43^, as a local scaffold that supports synapse reformation at muscle targets.

### Nidogen, integrin, and laminin form a tripartite mechanism to guide regenerating axons

Our results indicate that NID-1 functions together with laminin and integrin to guide regenerating axons. Laminin is a major component of the basement membrane and a known interactor of Nidogen^47^, while integrins are cell-surface receptors that link extracellular matrix cues to intracellular signaling pathways^57^. Integrins have well-established roles in both axon development and regeneration. In *C. elegans*, integrins are required for GABAergic axon guidance during development^58^, and promote axon regeneration across multiple organisms^59,60^. Here, we find that loss of integrin function (*ina-1(gm144)*) disrupts the ability of regenerating cholinergic axons to extend alongside PVD dendrites, phenocopying loss of *nid-1* (Figures 5D, 6B). Cholinergic motor neurons express higher levels of integrin than GABAergic neurons^44^, and regenerating GABAergic axons colocalize with PVD dendrites significantly less frequently than cholinergic axons (Figure 5A), consistent with the hypothesis that integrin levels influence trajectory choice of specific types of motor neurons. The finding that overexpression of either INA-1 or PAT-3 in GABAergic neurons redirects their regenerating axons along PVD dendrite trajectories (Figure 5G) confirms that integrin levels are a key determinant of trajectory choice. The subsequent findings that ectopic guidance provided by integrin overexpression in GABA neurons depends on NID-1 (Figure 6A), and that loss of *ina-1* and *nid-1* do not show additive effects on colocalization (Figure 5F), indicate integrin and NID-1 function together to direct regenerating axons. Loss of laminin (by RNAi knockdown of *lam-1*, *lam-2*, or *epi-1*) also disrupts the ability of regenerating cholinergic axons to extend alongside PVD dendrites (Figure 5I). Consistent with a previous finding that nidogen and laminin function together to form stable complexes along mechanosensory neurons^61^, the lack of additive phenotype between *nid-1*, *lam-1*, *lam-2* and *epi-1* indicates they function together to guide regenerating axons.

Taken together, these results reveal that laminin, nidogen, and integrin form a tripartite pathway that directs regenerating cholinergic motor axons alongside the PVD mechanosensory neuron to restore functional connectivity after injury. The preferential use of this pathway by cholinergic rather than GABAergic axons reflects their higher integrin expression^44^, revealing how cell-type-specific differences in ECM receptor levels can shape regenerative outcomes in a shared environment. These findings establish a framework for understanding how molecular cues in the mature nervous system direct regenerating axons to their targets after injury.

## Acknowledgments

We thank all members of the Byrne laboratory, Dr. Mark Alkema, Dr. Yang Xiang, and Dr. Michael Francis for their experimental suggestions, insight, and critical reading of the manuscript. We thank the Caenorhabditis Genetics Center (CGC), which is funded by the NIH Office of Research Infrastructure Program (P40 OD010440), for providing strains and WormBase. We thank the Alkema (UMass Chan Medical School) and Francis (UMass Chan Medical School) laboratories for plasmids and strains. **Funding:** National Institute of Neurological Disease and Stroke, R01NS110936; National Institute of Neurological Disease and Stroke, F31NS139608.

## Author contributions

E.M. contributed to investigation, interpretation, and manuscript preparation. W.H. contributed to conceptualization and interpretation. A.B.B contributed to conceptualization, interpretation, and manuscript preparation.

## Data and materials availability

All data is available in the main text or the supplementary materials. Strains will be made available upon request.

## Material and Methods

### Strains

Animals were maintained at 20°C on NGM plates containing OP50 *E. coli* according to standard methods. ABC120 *ufIs43[acr-2::mCherry]; wdIs51[F49H12.4::GFP + unc-119(+)]*, ABC537 *ufIs43[acr-2::mCherry]; wdIs51[F49H12.4::GFP + unc-119(+)]; [ser-2prom3::deg-3-N293I],* ABC725 *wdIs51[F49H12.4::GFP + unc-119(+)]; ufIs43[acr-2::mCherry];nid-1(cg119),* ABC755 *wdIs51[F49H12.4::GFP + unc-119(+)]; ufIs43[acr-2::mCherry]; nid-1(cg118)*, ABC723 *wdIs51[F49H12.4::GFP + unc-119(+)]; ufIs43[acr-2::mCherry]; nid-1(cg119); bamEx335[dpy-7p::nid-1A]*, ABC707 *wdIs51[F49H12.4::GFP + unc-119(+)]; ufIs43[acr-2::mCherry]; nid-1(cg119); bamEx336[dpy-7p::nid-1A]*, ABC728 *wdIs51[F49H12.4::GFP + unc-119(+)]; ufIs43[acr-2::mCherry]; nid-1(cg119); bamEx337[myo-3p::nid-1A]*, ABC722 *wdIs51[F49H12.4::GFP + unc-119(+)]; ufIs43[acr-2::mCherry]; nid-1(cg119); bamEx338[myo-3p::nid-1A]*, ABC807 *wdIs51[F49H12.4::GFP + unc-119(+)]; ufIs43[acr-2::mCherry]; nid-1(cg119); bamEx333[unc-17::nid-1A]*, ABC753 *wdIs51[F49H12.4::GFP + unc-119(+)]; ufIs43[acr-2::mCherry]; nid-1(cg119); bamEx334[unc-17::nid-1A]*, ABC700 *ufIs43[acr-2::mCherry]; ufIs233[unc-129p::Dendra2::RAB-3]*, ABC704 *ufIs43[acr-2::mCherry]; ufIs233[unc-129p::Dendra2::RAB-3]; nid-1(cg119)*, ABC741 *ufIs43[acr-2::mCherry]; ufIs233[unc-129p::Dendra2::RAB-3]; nid-1(cg118)*, ABC793 *ufIs43[acr-2::mCherry]; ufIs233[unc-129p::Dendra2::RAB-3]; nid-1(cg119); bamEx337[myo-3p::nid-1A]*, ABC746 *ufIs43[acr-2::mCherry]; ufIs233[unc-129p::Dendra2::RAB-3]; nid-1(cg119); bamEx338[myo-3p::nid-1A]*, ABC726 *ufIs43 [acr-2::mCherry]; ufIs233[unc-129p::Dendra2::RAB-3]; nid-1(cg119); bamEx335[dpy-7p::nid-1A::unc54],* ABC718 *ufIs43[acr-2::mCherry]; ufIs233[unc-129p::Dendra2::RAB-3]; nid-1(cg119); bamEx336[dpy-7p::nid-1A],* ABC706 *wdIs51[F49H12.4::GFP + unc-119(+)]; wpIs40 [unc-47p::mCherry],* ABC731 *wdIs51[F49H12.4::GFP + unc-119(+)]; ufIs43[acr-2::mCherry]; ina-1(gm144),* ABC803 *wdIs51[F49H12.4::GFP + unc-119(+)]; ufIs43[acr-2::mCherry]; bamEx340[unc-47::ina-1],* ABC737 *wdIs51[F49H12.4::GFP + unc-119(+)]; ufIs43[acr-2::mCherry]; bamEx341[unc-47::ina-1],* ABC739 *wdIs51[F49H12.4::GFP + unc-119(+)]; ufIs43[acr-2::mCherry]; bamEx342[unc-47::pat-3],* ABC740 *wdIs51[F49H12.4::GFP + unc-119(+)]; ufIs43[acr-2::mCherry]; bamEx342[unc-47::pat-3]*.

### Axotomy and Microscopy

Axotomy experiments were performed as previously described^62^. Larval stage 4 (L4) worms were mounted on 3% agarose pads and immobilized with 1:12 polystyrene microbeads (Polysciences) diluted in water. Single injuries were made at the midline of the axon commissure and 1-2 cholinergic axons or 1-3 GABAergic axons were cut per worm. For axotomy experiments with PVD labeled strains, a single cut was performed on the motor axons avoiding damage to the PVD dendrites. Axon regeneration was quantified 24 hr after injury unless otherwise stated by placing animals immobilized with 300 nM sodium azide on a 3% agarose pad. Animals were imaged with a Nikon Eclipse 80i microscope, 100x Plan Apo VC lens (1.4 NA), Andor Zyla sCMOS camera and a Leica EL6000 light source with NIS-Elements AR5.02.00 software, or a Perkin Elmer UltraVIEW VoX confocal imaging system mounted on a Zeiss AXIO (imager.M2) microscope and Volocity 6.3 software. To control day-to-day and instrument-to-instrument variability, experiments were carried out with same day controls on one axotomy rig.

### Molecular biology

Tissue-specific rescue constructs were generated by amplifying *nid-1A* from cDNA. Plasmids expressing NID-1A under *unc-17* cholinergic neuronal promoter *pEM1[unc-17::nid-1A::let854]*, *myo-3* body wall muscle promoter *pEM2[myo-3p::nid-1A::unc54]*, and *dpy-7* hypodermal promoter *pEM3[dpy-7p::nid-1A::unc54]* were generated by Gibson cloning and confirmed by whole plasmid sequencing. Plasmids were injected into ABC725 *wdIs51[F49H12.4::GFP + unc-119(+)]; ufIs43 [acr-2::mCherry]; nid-1(cg119)*, which is a NID-1 null mutant with dual labeled cholinergic motor neurons and the PVD neurons at a final concentration of 15ng/µl.

Plasmids expressing *ina-1 (pDO117 [punc-47::ina-1])* and *pat-3 (pAP86 [unc-47::pat-3])* under the ***unc-47*** GABAergic neuron promoter were injected into ABC706 *wdIs51[F49H12.4::GFP + unc-119(+)]; wpIs40 [unc-47p::mCherry]* at a final concentration of 2ng/µl. Plasmids *pDO117* and *pAP86* were a gift from the Francis Lab at UMass Chan Medical School.

### Aldicarb assay

Strains were scored in parallel, with the scorer blinded to the genotype and condition. Young adult animals (24 h after L4) were selected (>5 per trial for at least 3 trials) and transferred to a plate containing 1 mM aldicarb and nematode growth medium. Movement was assessed every 10 min for 180 minutes. Animals that displayed no movement when prodded (paralyzed) were noted. The percentage of paralyzed animals was calculated at each timepoint.

### RNAi

RNAi clones (V-8J05, IV-2P15, X-3B20, and IV-5J23) from the *C. elegans* RNAi feeding library were used to knock down *nid-1, lam-1, lam-2,* and *epi-1,* respectively. RNAi strains were cultured overnight in LB with 25 μg/mL carbenicillin and 12.5 μg/mL tetracycline. The resulting culture was diluted 1:100 in LB + 25 μg/mL carbenicillin for 6h (Jedrusik & Schulze, 2004) and 400µl RNAi culture was seeded on NGM plates containing 25 μg/mL carbenicillin and 1 mM IPTG. Strains fed L4440 (empty vector) and *unc-22* (clone IV-6K06, knock down causes a twitching phenotype) RNAi were used as negative and positive controls in parallel. Animals were fed RNAi for 2 generations and axotomized according to the axotomy protocol. Animals were recovered to RNAi plates after axotomy.

### Dendra2::RAB-3 Synapse Reformation

The starting strain *ufIs233[unc-129p::Dendra2::RAB-3]* was a gift from the Francis Lab at UMass Chan Medical School. L4 animals from wild-type, *nid-1(cg118)*, *nid-1(cg119)*, and *nid-1A* rescue strains expressing Dendra2::RAB-3 were axotomized according to the axotomy protocol described above. Dorsal RAB-3 puncta of the injured DA/DB axons were photoconverted using a 405 nm laser at 800 ms for 45 s^35^. Photoconverted animals were rescued onto *E. coli* OP50-seeded plates for 24 hours before quantifying green-fluorescent RAB-3 puncta. A non-parametric Kruskal-Wallis test was used for statistical analysis.

### Synaptic puncta analysis

Background fluorescence was subtracted by calculating the average intensity in a region outside the ROI. A set ROI of 52 µm in length was used for all analysis. Puncta within an ROI were quantified using set threshold of 65-255 and the analyze particles function of ImageJ was used to quantify any particle >4 pixels. For quantification of fully regenerated cholinergic neuron synapses, the ROI was defined as 26 µm anterior and posterior to where the regenerated axons join the dorsal nerve cord. For quantification of the uninjured cholinergic neuron synapses the ROI was defined as 26 µm anterior and posterior to where the axons join the dorsal nerve cord.

---

**Supplemental Figure 1.**
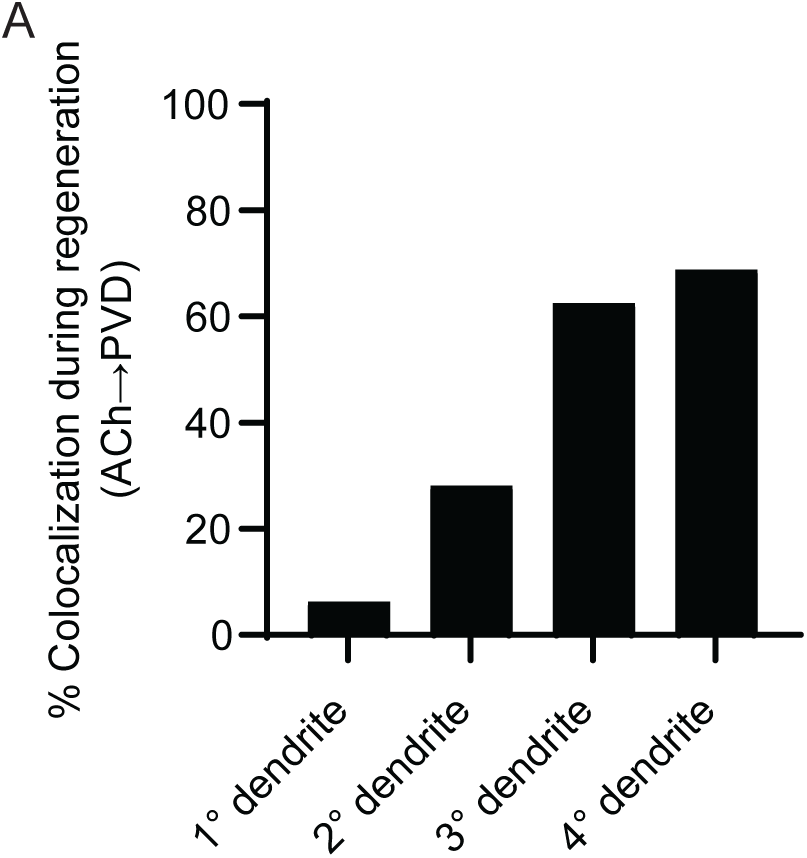
Regenerating cholinergic axons colocalize with multiple levels of PVD dendritic branches. **(A)** Regenerating cholinergic axons colocalize with all levels of the PVD’s dendritic branches, and most frequently with 3° and 4° branches (N=6,28,62,69). Individual axons are represented in multiple bars if they colocalize with more than one level of branch.

**Supplemental Figure 2.**
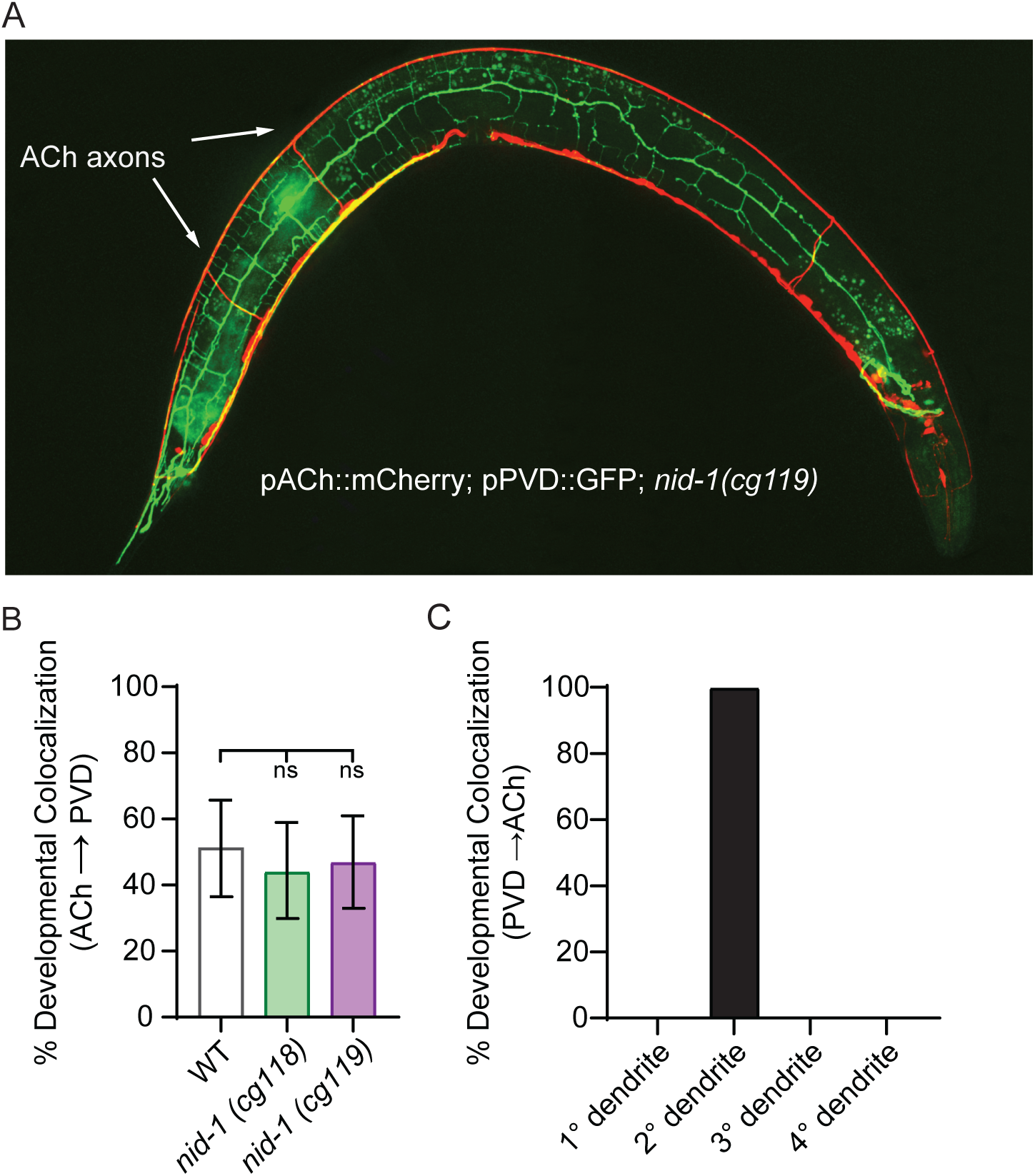
Loss of *nid-1* does not affect developmental colocalization between ACh axons and PVD dendrites. **(A)** Representative image of P*acr-2*::mCherry; P*pvd*::GFP; *nid-1(cg119)* animal showing ACh commissures (red) and PVD dendrites (green). **(B)** The proportion of ACh axons that colocalize with PVD dendrites during development is not significantly different in *nid-1(cg118)* or *nid-1(cg119)* mutants compared to wild-type. **(C)** During development, when colocalization between a PVD dendrite and an ACh axon is observed, it occurs exclusively along 2° dendritic branches, as previously reported (N=0,25,0,0)^30^.

**Supplemental Figure 3.**
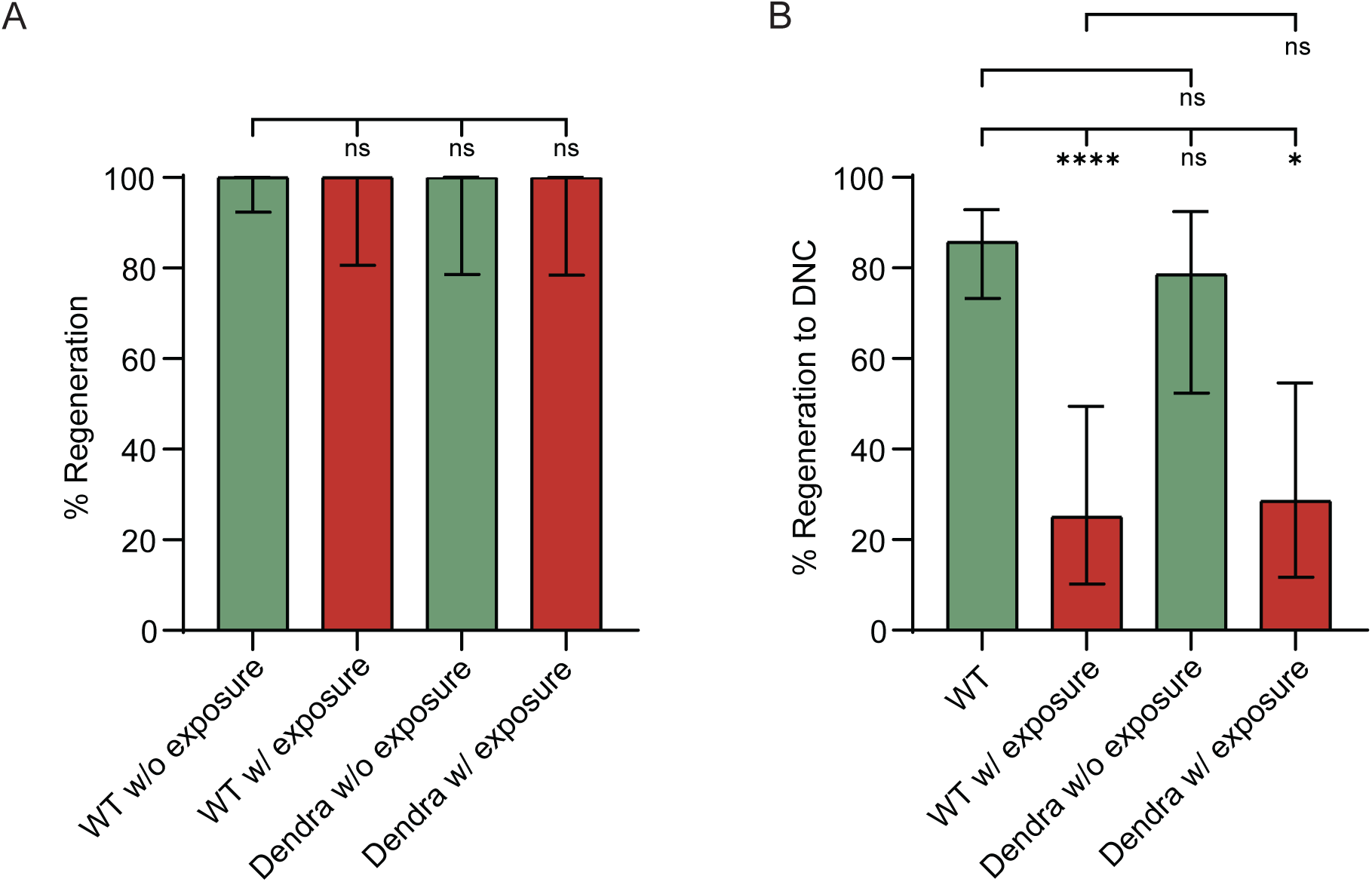
Exposure to 405nm light does not disrupt the initiation of axon regeneration but negatively influences axon extension. **(A)** The proportion of axons that initiate regeneration does not differ between animals with or without exposure to 405nm photoconversion light, regardless of Dendra2::RAB-3 expression (N=49,16,14,14). **(B)** Exposure to 405nm light, but not Dendra2::RAB-3 expression alone, decreases the proportion of regenerating axons that reach the dorsal cord (N=49,16,14,14). Significance determined by Fisher’s exact test and is indicated by *p≤0.05 and ****p≤0.0001. Error bars represent 95% confidence intervals.

**Supplemental Figure 4.**
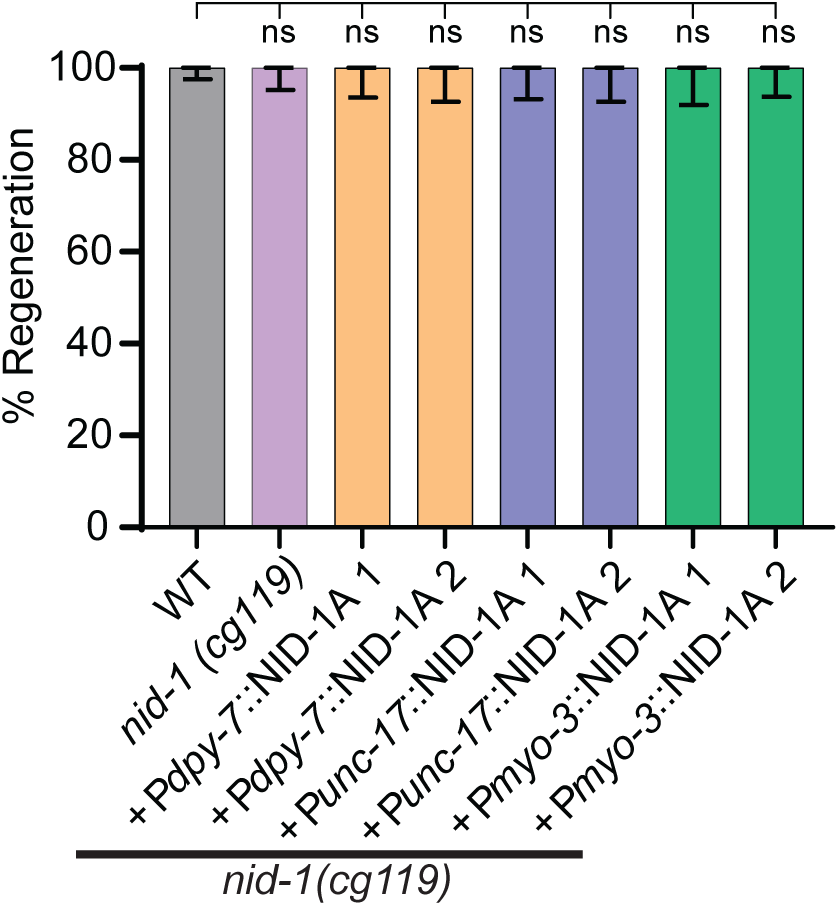
Expressing NID-1 in the body wall muscles or hypodermis does not alter the initiation of regeneration. The proportion of axons that initiate regeneration is not significantly altered in any of the tissue-specific rescue genotypes compared to *nid-1(cg119)* (N=150,77,56,48,53,48,44,57). Error bars represent 95% confidence intervals.

**Supplemental Figure 5.**
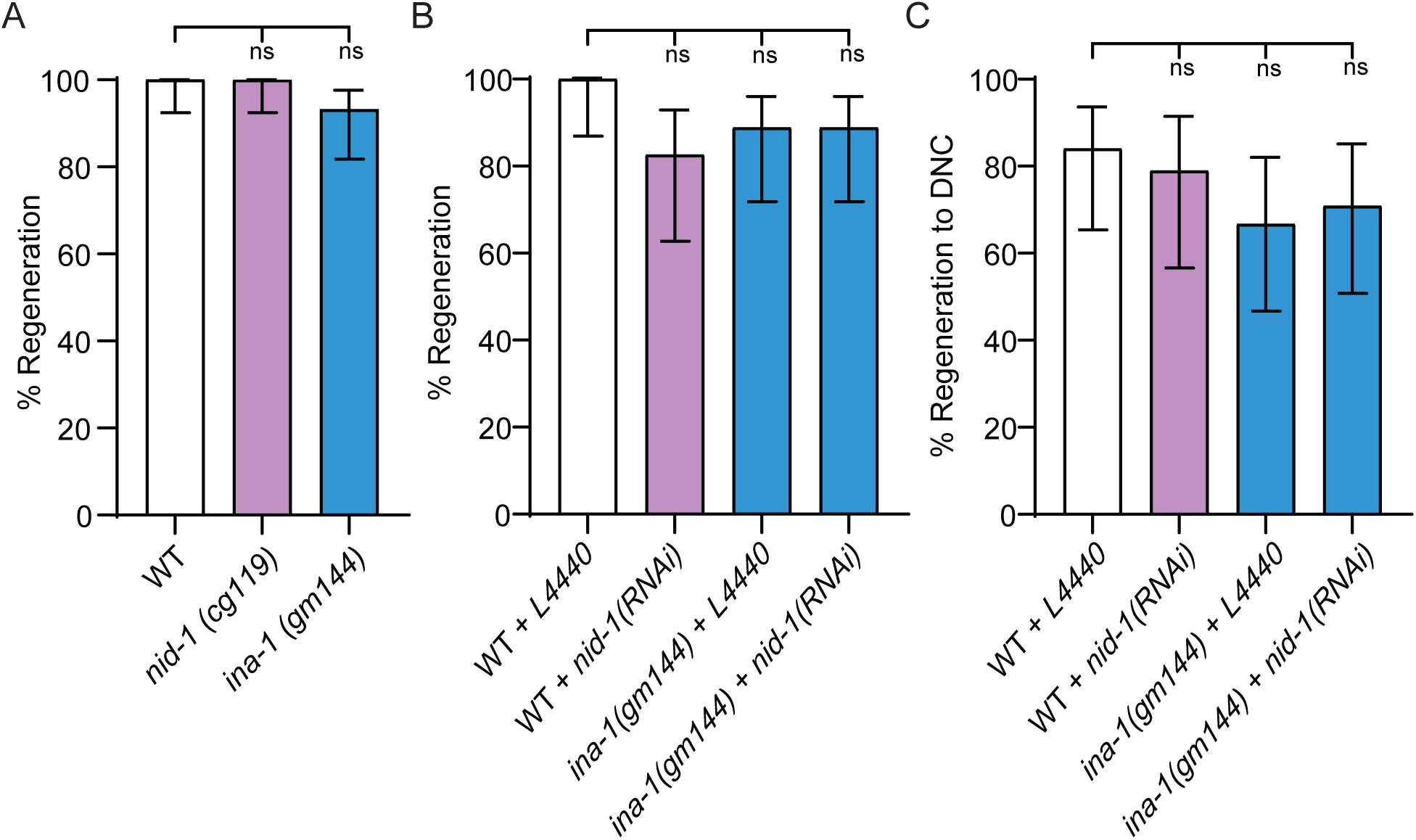
Loss of *nid-1* or *ina-1* does not alter the initiation of regeneration. **(A)** The proportion of injured axons that initiate regeneration in *nid-1(cg119)* and *ina-1(gm144)* mutants does not differ significantly from wild-type (N=47,47,44). **(B)** The proportion of injured axons that initiate regeneration in wild-type animals fed *nid-1* RNAi, *ina-1(gm144)* animals fed empty RNAi vector (L4440), or *ina-1(gm144)* animals fed *nid-1* RNAi does not differ significantly from wild-type animals fed empty RNAi vector (N=25,23,27,27). **(C)** The proportion of regenerating axons that reach the dorsal cord in wild-type animals fed *nid-1* RNAi, *ina-1(gm144)* animals fed empty RNAi vector (L4440), or *ina-1(gm144)* animals fed *nid-1* RNAi does not differ significantly from wild-type animals fed empty RNAi vector (N=25,19,24,24).

**Supplemental Figure 6.**
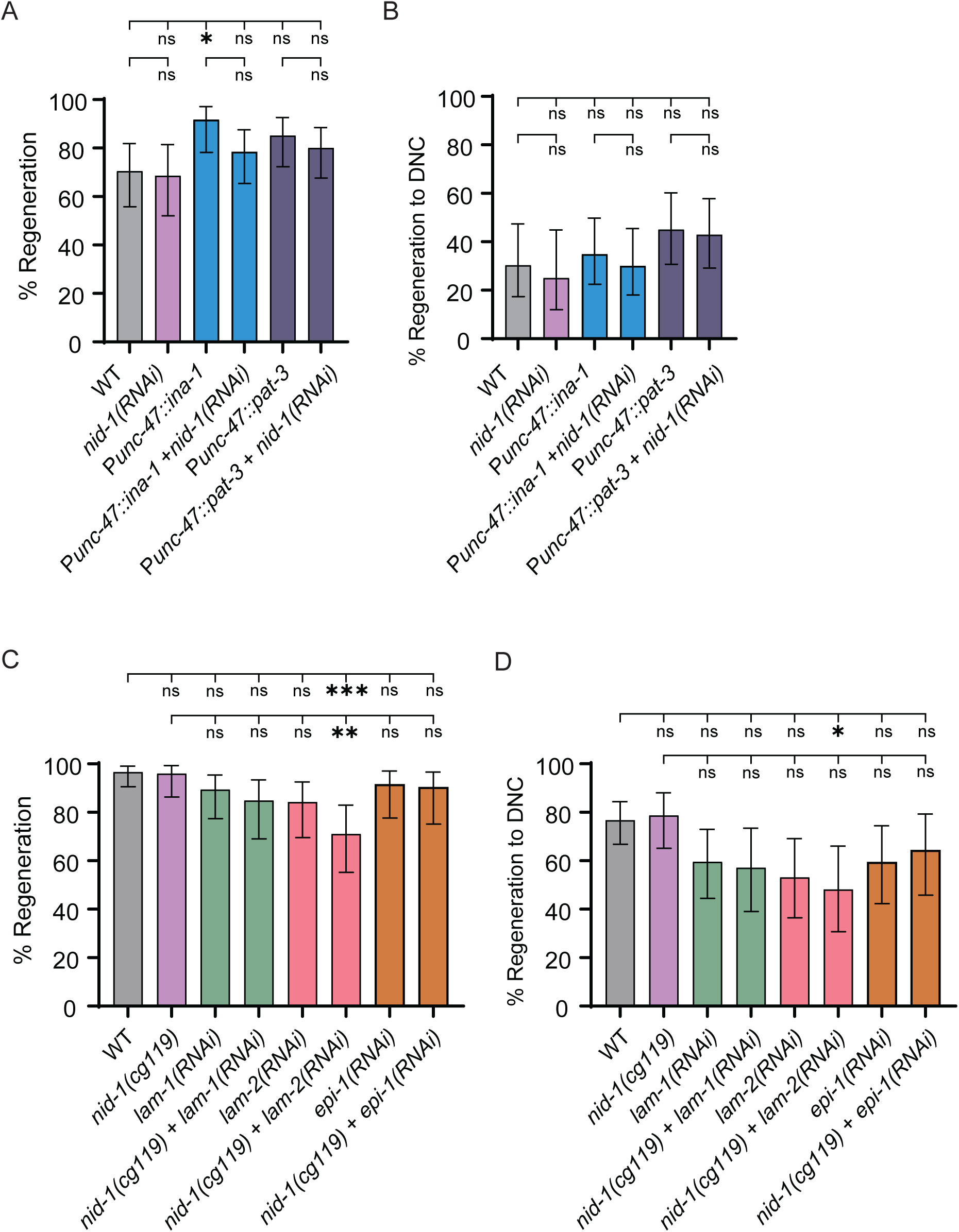
Regeneration frequency, regeneration initiation and dorsal cord extension of nidogen, integrin, and laminin mutants. **(A)** *nid-1* RNAi does not affect the proportion of GABAergic axons that initiate regeneration in wild-type animals, or in animals that overexpress *ina-1* or *pat-3* in GABAergic neurons (P*unc-47*::*ina-1* or P*unc-47*::*pat-3*) (N=44,35,36,51,47,55). **(B)** *nid-1* RNAi does not affect the proportion of GABAergic axons that reach the dorsal nerve cord (N=33,24,43,40,40,42). **(C)** LAM-2 RNAi significantly decreases the proportion of ACh axons that initiate regeneration, while *lam-1* and *epi-1* RNAi do not (N=89,49,47,33,38,38,35,31). **(D)** LAM-2 RNAi significantly decreases the proportion of ACh axons that reach the dorsal cord, while *lam-1* and *epi-1* RNAi do not (N=86,47,42,28,32,27,32,28). Significance determined by Fisher’s exact test and is indicated by *p≤0.05, **p≤0.01, and ***p≤0.001. Error bars represent 95% confidence intervals.

